## Supplemental Figures and Tables for "Immigrant birds learn from socially observed differences in payoffs when their environment changes"

### Supplementary tables and figures for manuscript "Immigrant birds learn from socially observed differences in payoffs when their environment changes"

#### Figures


- **Figure S1**: Results from preference tests
- **Figure S2**: Daily preferences of residents after immigration.
- **Figure S3**: WAIC model comparison
- **Figure S4**: Individual point estimates for key parameters
- **Figure S5**: New associate bias estimates from model SL3
- **Figure S6**: Tutor bias estimates for immigrants


#### Tables


- **Table S1**: Relative value of food items.
- **Table S2**: Non-tutor birds preferred the seeded solution.
- **Table S3**: How did immigrant preferences change after the immigration event?
- **Table S4**: Summary of model estimates, individual learning only (IL)
- **Table S5**: Summary of model estimates, SL1
- **Table S6**: Summary of model estimates, SL2
- **Table S7**: Summary of model estimates, SL3

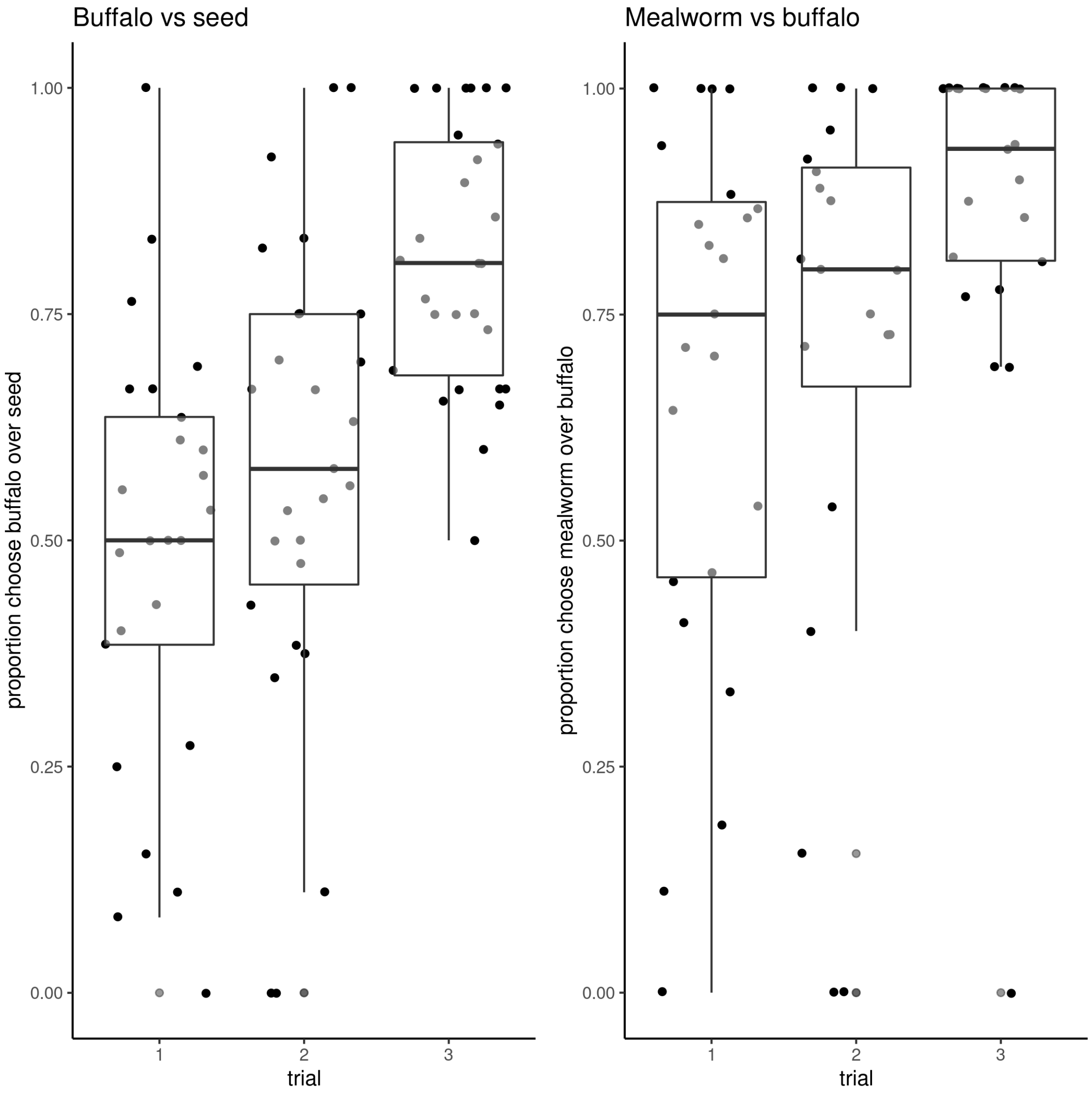


Figure S1: **Results from preference tests.** Proportion of choices (y-axis) by trial (x-axis), where each dot represents an individual bird. Birds preferred buffalo worms to seed, and mealworms to buffalo mealworms, and these preferences strengthened over time. The data underlying this figure can be found in our data and code repository `https://doi.org/10.17617/3.FXC12W`.

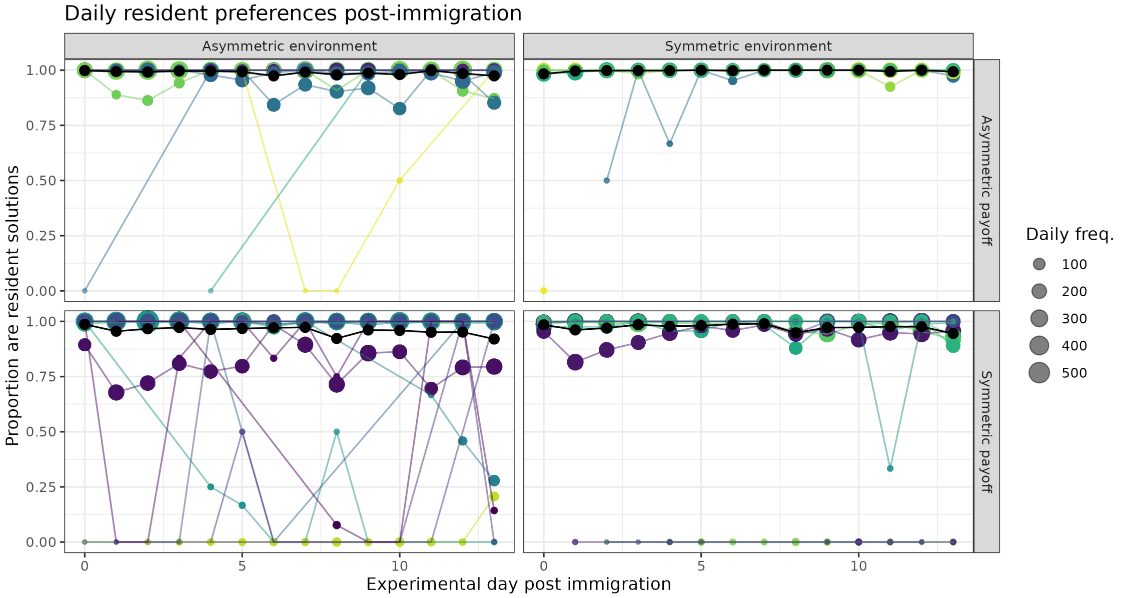


Figure S2: **Daily preferences of residents after immigration.** Proportion of immigrant solutions which were resident solutions (y-axis) over experimental day (x-axis). Colored lines are individual immigrants, with the daily solving frequency indicated (size). A handful of residents in the $P_s$ conditions adopted the immigrant preference, however they had either learned to use the puzzle after immigration, or had very low solving rates. The data underlying this figure can be found in our data and code repository`https://doi.org/10.17617/3.FXC12W`.

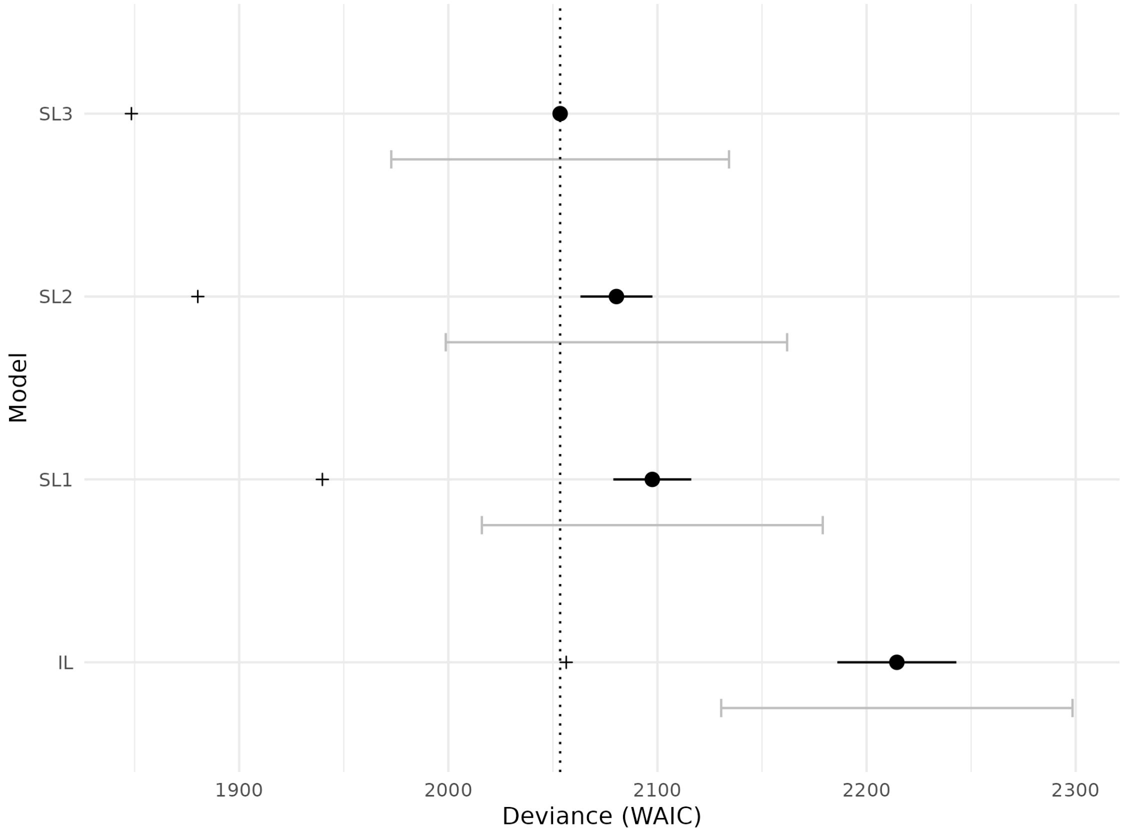


| | WAIC | SE | dWAIC | dSE | pWAIC | weight |
|---|---|---|---|---|---|---|
| *SL3* | 2053.46 | 80.76 | 0 | NA | 102.51 | 1 |
| *SL2* | 2080.38 | 81.62 | 26.92 | 17.22 | 100.09 | 0 |
| *SL1* | 2097.53 | 81.5 | 44.07 | 18.64 | 78.88 | 0 |
| *IL* | 2214.47 | 84 | 161.01 | 28.46 | 79.05 | 0 |

Figure S3: **WAIC model comparison**. Plot and summary table for WAIC comparison of dynamic learning models. Black dots indicate out-of-sample deviance, + symbols indicate in-sample deviance. Grey bars indicate standard error, and black bars are the difference in standard errors. Vertical dotted line is the WAIC of the highest ranked model. Models with social learning components were better fit than individual learning alone, and the highest ranked model estimated payoff-biased social learning, a new associate bias, and a sloped change to the sensitivity to social information. The data underlying this figure can be found in our data and code repository`https://doi.org/10.17617/3.FXC12W`.

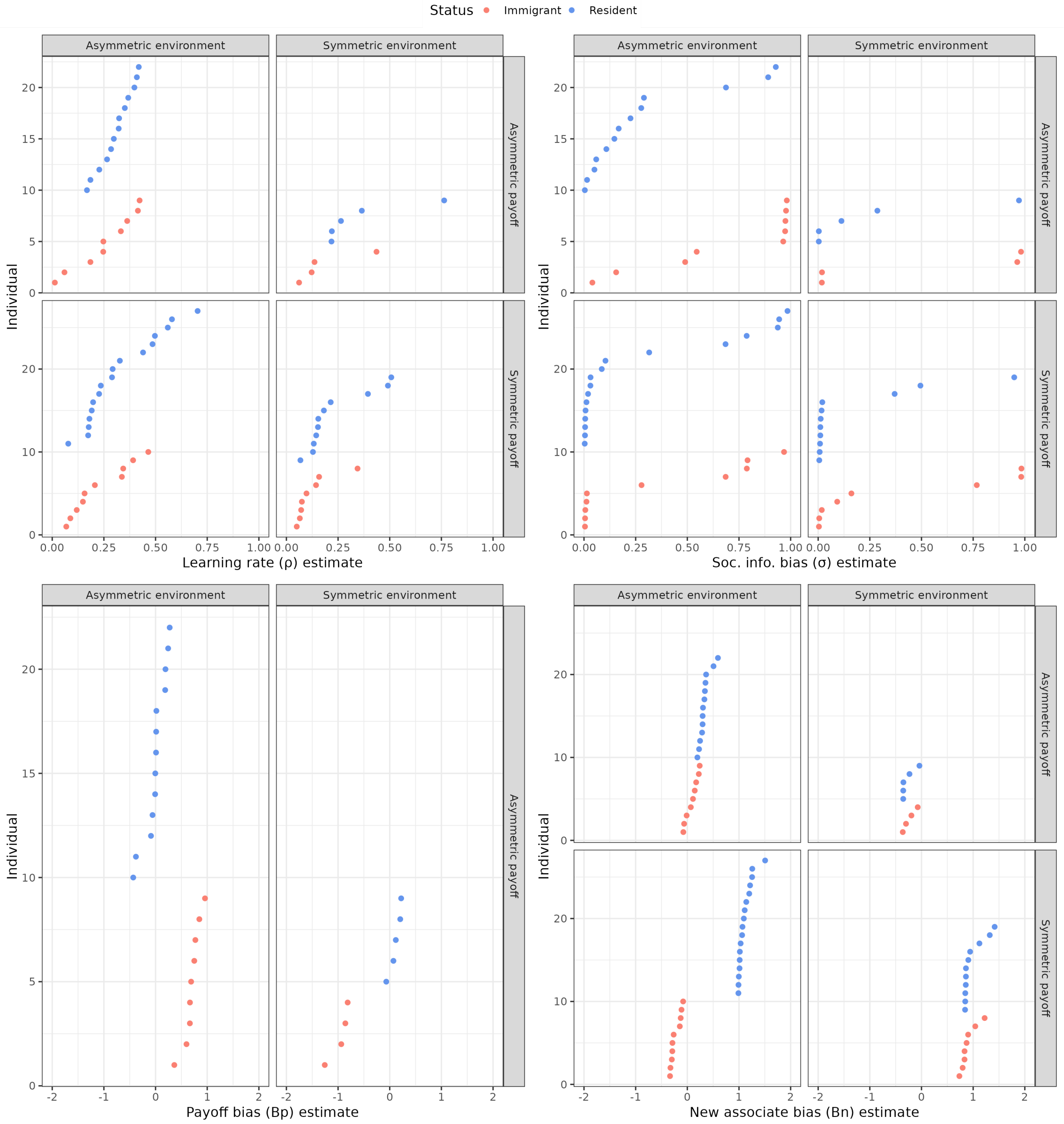


Figure S4: **Individual point estimates for key parameters**. Point estimates of individual birds from Bayesian learning model SL3 (immigrant status as color). Point estimates shown for learning rate ($\rho$), social information bias ($\sigma$), payoff bias ($\beta_p$), new associate bias ($\beta_n$). The data underlying this figure can be found in our data and code repository`https://doi.org/10.17617/3.FXC12W`.

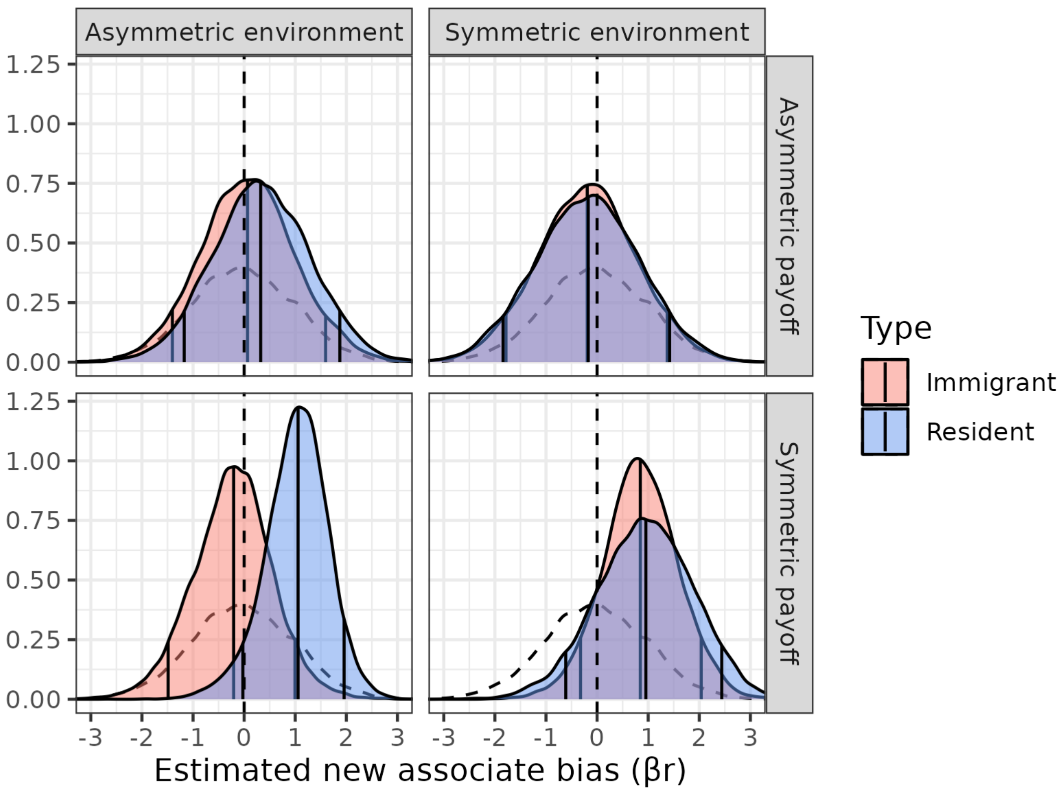


Figure S5: **New associate bias estimates from model SL3**. Estimates above zero indicate that birds were disproportionately influenced by observations of solves by new associates. Estimates below zero indicate that birds ignored observations of solves by new associates. The data underlying this figure can be found in our data and code repository`https://doi.org/10.17617/3.FXC12W`.

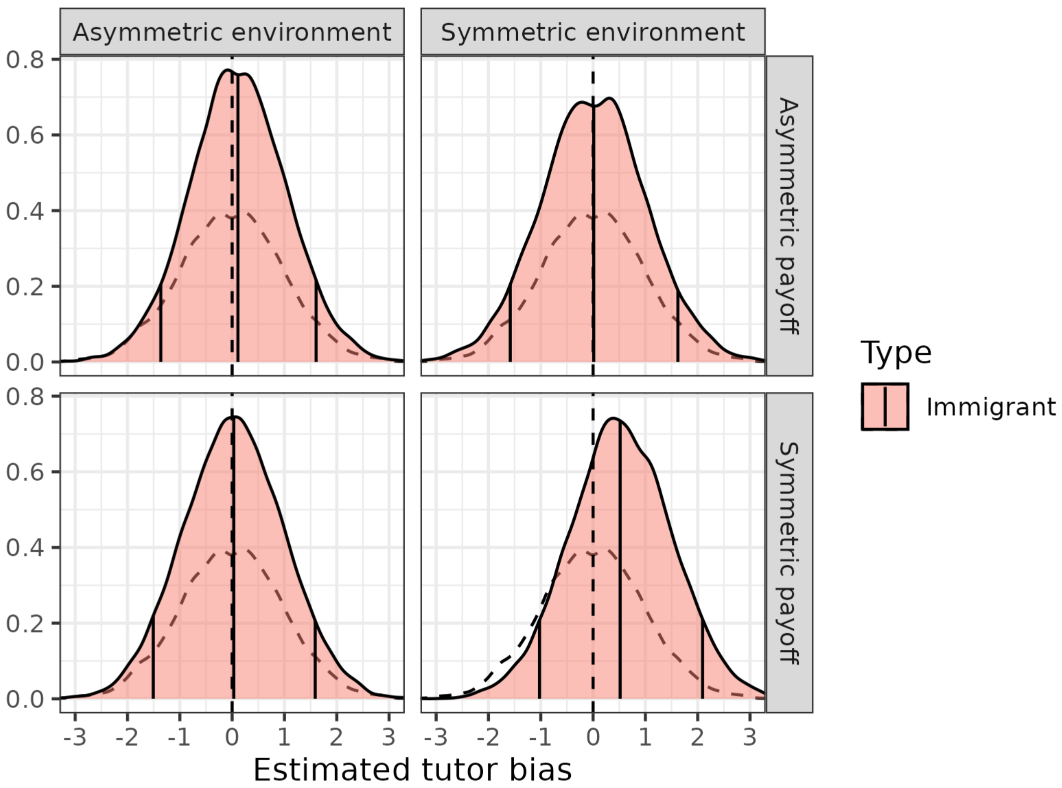


Figure S6: **Tutor bias estimates for immigrants**. We fit an additional model, similar to SL3, except that rather than information about whether a solution was produced by a new associate, we included information about whether a solution was produced by a tutor. This was done to determine whether immigrants were disproportionately influenced by the productions of tutors in their new populations. We found no evidence to support this in any condition. The data underlying this figure can be found in our data and code repository`https://doi.org/10.17617/3.FXC12W`.

Table S1: **Relative value of food items.** Value estimates from Bayesian inverse reinforcement learning model. Reported values are mean (89% HDPI). The data underlying this table can be found in our data and code repositoryhttps://doi.org/10.17617/3.FXC12W.

| parameter | value | n. eff. | Rhat |
|---|---|---|---|
| reward_seeds | 0.35 (0.02,1.02) | 1275.45 | 1.00 |
| reward_buffalo | 1.72 (1.33,2.39) | 1294.75 | 1.00 |
| reward_mealworms | 3.66 (3.12,4.4) | 1444.63 | 1.00 |

Table S2: **Non-tutor birds preferred the seeded solution.** Linear mixed model where the response variable was the proportion of solutions by non-tutors during the diffusion period that were of the seeded side. There were no significant differences between age, sex, immigrants, residents or conditions. Year was included as a random intercept. The data underlying this table can be found in our data and code repository`https://doi.org/10.17617/3.FXC12W`.

| | *Dependent variable:* |
|---|---|
| | Prop. seeded soln. |
| Age (adult) | −0.056 (0.034) |
| | t = −1.628 |
| | p = 0.104 |
| Sex (male) | 0.054 (0.033) |
| | t = 1.614 |
| | p = 0.107 |
| Resident | 0.048 (0.063) |
| | t = 0.764 |
| | p = 0.446 |
| Payoff condition (symmetric) | −0.070 (0.062) |
| | t = −1.120 |
| | p = 0.263 |
| Environment condition (symmetric) | 0.014 (0.070) |
| | t = 0.195 |
| | p = 0.846 |
| Resident:Payoff condition | −0.070 (0.080) |
| | t = −0.866 |
| | p = 0.387 |
| Resident:Environment condition | −0.015 (0.102) |
| | t = −0.150 |
| | p = 0.881 |
| Payoff condition:Environment condition | 0.052 (0.093) |
| | t = 0.563 |
| | p = 0.574 |
| Resident:Payoff condition:Environment condition | 0.021 (0.130) |
| | t = 0.163 |
| | p = 0.871 |
| Intercept | 0.995 (0.067) |
| | t = 14.842 |
| | p = 0.000*** |
| Observations | 71 |
| Log Likelihood | 27.199 |
| Akaike Inf. Crit. | −30.397 |
| Bayesian Inf. Crit. | −3.245 |
| *Note:* | *p<0.1; **p<0.05; ***p<0.01 |

Table S3: **How did immigrant preferences change after the immigration event?** Logistic GLMM where the probability of producing the resident solve was predicted by normalized experience solving the puzzle pre-immigration (0 representing the mean number of solutions produced of the resident side of an immigrants original population), and the interaction of days since immigration (starting at 0), payoff condition and environmental condition. The intercept can be interpreted as the probability of producing the resident solution of a bird with average experience of solving on the day immediately following immigration. The data underlying this table can be found in our data and code repositoryhttps://doi.org/10.17617/3.FXC12W.

| | *Dependent variable:* |
|---|---|
| | Produce resident soln. |
| Normalized experience | −1.036 (0.674) |
| | t = −1.538 |
| | p = 0.125 |
| Day since immigration | 0.223 (0.008) |
| | t = 26.788 |
| | p = 0.000*** |
| Environment condition (symmetric) | −2.156 (2.367) |
| | t = −0.911 |
| | p = 0.363 |
| Payoff condition (symmetric) | −6.550 (1.721) |
| | t = −3.805 |
| | p = 0.0002*** |
| Day:Environment condition | 0.431 (0.023) |
| | t = 18.413 |
| | p = 0.000*** |
| Day:Payoff condition | −0.032 (0.013) |
| | t = −2.427 |
| | p = 0.016** |
| Payoff condition:Environment condition | 3.956 (2.948) |
| | t = 1.342 |
| | p = 0.180 |
| Day:Environment condition:Payoff condition | −0.765 (0.034) |
| | t = −22.450 |
| | p = 0.000*** |
| Intercept | 2.153 (1.232) |
| | t = 1.747 |
| | p = 0.081* |
| Observations | 40,363 |
| Log Likelihood | −8,000.502 |
| Akaike Inf. Crit. | 16,023.000 |
| Bayesian Inf. Crit. | 16,117.670 |
| *Note:* | *p<0.1; **p<0.05; ***p<0.01 |

Table S4: **Summary of model estimates, individual learning only (IL)**. Table contains mean values (89% HDPI), number of effective samples, and Gelman's Rhat. Parameter estimates and SD of varying effects across individuals are reported. The data underlying this table can be found in our data and code repositoryhttps://doi.org/10.17617/3.FXC12W.

| condition | value | n. eff. | Rhat |
|---|---|---|---|
| **Recent information bias** $\rho$ | | | |
| $E_a, P_a$ res. | 0.32 (0.14,0.58) | 7828.97 | 1.00 |
| $E_a, P_a$ immi. | 0.29 (0.14,0.52) | 5312.50 | 1.00 |
| $E_a, P_s$ res. | 0.28 (0.14,0.5) | 7424.18 | 1.00 |
| $E_a, P_s$ immi. | 0.22 (0.09,0.45) | 6048.09 | 1.00 |
| $E_s, P_a$ res. | 0.33 (0.12,0.64) | 10894.80 | 1.00 |
| $E_s, P_a$ immi. | 0.29 (0.11,0.6) | 6167.35 | 1.00 |
| $E_s, P_s$ res. | 0.27 (0.12,0.5) | 8152.20 | 1.00 |
| $E_s, P_s$ immi. | 0.16 (0.06,0.38) | 7480.34 | 1.00 |
| **Behavioral conservatism** $\alpha$ | | | |
| | 1.29 (1.05,1.59) | 3693.88 | 1.00 |
| **SD individals** | | | |
| rho (logit scale) | 2.13 (1.49,2.93) | 3772.24 | 1.00 |
| alpha (log scale) | 0.97 (0.76,1.23) | 4401.25 | 1.00 |

Table S5: **Summary of model estimates, SL1.** Table contains mean values (89% HDPI), number of effective samples, and Gelman's Rhat. Parameter estimates and SD of varying effects across individuals are reported. The data underlying this table can be found in our data and code repositoryhttps://doi.org/10.17617/3.FXC12W.

| condition | value | n. eff. | Rhat |
|---|---|---|---|
| **Recent information bias** $\rho$ | | | |
| $E_a, P_a$ res. | 0.43 (0.19,0.71) | 9693.42 | 1.00 |
| $E_a, P_a$ immi. | 0.3 (0.13,0.58) | 4360.72 | 1.00 |
| $E_a, P_s$ res. | 0.47 (0.24,0.71) | 5511.47 | 1.00 |
| $E_a, P_s$ immi. | 0.33 (0.14,0.6) | 4454.35 | 1.00 |
| $E_s, P_a$ res. | 0.46 (0.19,0.75) | 11384.45 | 1.00 |
| $E_s, P_a$ immi. | 0.32 (0.12,0.63) | 5468.66 | 1.00 |
| $E_s, P_s$ res. | 0.38 (0.17,0.65) | 5507.65 | 1.00 |
| $E_s, P_s$ immi. | 0.21 (0.07,0.5) | 4216.93 | 1.00 |
| **Social information bias** $\sigma$ | | | |
| $E_a, P_a$ res. | 0.24 (0.07,0.58) | 14778.65 | 1.00 |
| $E_a, P_a$ immi. | 0.27 (0.08,0.61) | 15717.76 | 1.00 |
| $E_a, P_s$ res. | 0.13 (0.03,0.38) | 11849.47 | 1.00 |
| $E_a, P_s$ immi. | 0.15 (0.04,0.44) | 14627.14 | 1.00 |
| $E_s, P_a$ res. | 0.24 (0.06,0.6) | 16923.20 | 1.00 |
| $E_s, P_a$ immi. | 0.28 (0.08,0.64) | 16501.87 | 1.00 |
| $E_s, P_s$ res. | 0.14 (0.03,0.42) | 14985.13 | 1.00 |
| $E_s, P_s$ immi. | 0.23 (0.06,0.58) | 15753.78 | 1.00 |
| **Behavioral conservatism** $\alpha$ | | | |
| | 1.45 (1.08,1.96) | 3477.45 | 1.00 |
| **SD individals** | | | |
| rho (logit scale) | 2.49 (1.67,3.47) | 3225.22 | 1.00 |
| sigma (logit scale) | 7.32 (5.19,10.01) | 6499.56 | 1.00 |
| alpha (log scale) | 0.97 (0.7,1.3) | 4030.65 | 1.00 |

Table S6: **Summary of model estimates, SL2.** Table contains mean values (89% HDPI), number of effective samples, and Gelman's Rhat. Parameter estimates and SD of varying effects across individuals are reported. The data underlying this table can be found in our data and code repository`https://doi.org/10.17617/3.FXC12W`.

| condition | value | n. eff. | Rhat |
|---|---|---|---|
| **Recent information bias $\rho$** | | | |
| $E_a, P_a$ res. | 0.4 (0.17,0.69) | 12083.94 | 1.00 |
| $E_a, P_a$ immi. | 0.26 (0.1,0.54) | 7302.48 | 1.00 |
| $E_a, P_s$ res. | 0.45 (0.23,0.71) | 8683.09 | 1.00 |
| $E_a, P_s$ immi. | 0.32 (0.13,0.59) | 6662.80 | 1.00 |
| $E_s, P_a$ res. | 0.46 (0.18,0.77) | 18116.73 | 1.00 |
| $E_s, P_a$ immi. | 0.33 (0.12,0.63) | 8183.72 | 1.00 |
| $E_s, P_s$ res. | 0.35 (0.14,0.63) | 9591.48 | 1.00 |
| $E_s, P_s$ immi. | 0.22 (0.07,0.51) | 7797.58 | 1.00 |
| **Social information bias $\sigma$** | | | |
| $E_a, P_a$ res. | 0.25 (0.07,0.6) | 18884.55 | 1.00 |
| $E_a, P_a$ immi. | 0.3 (0.09,0.65) | 20337.34 | 1.00 |
| $E_a, P_s$ res. | 0.12 (0.03,0.37) | 16612.20 | 1.00 |
| $E_a, P_s$ immi. | 0.16 (0.04,0.47) | 19557.11 | 1.00 |
| $E_s, P_a$ res. | 0.25 (0.07,0.61) | 25372.26 | 1.00 |
| $E_s, P_a$ immi. | 0.28 (0.08,0.64) | 23219.85 | 1.00 |
| $E_s, P_s$ res. | 0.16 (0.04,0.48) | 11654.37 | 1.00 |
| $E_s, P_s$ immi. | 0.23 (0.06,0.58) | 22053.23 | 1.00 |
| **Payoff bias $\beta_p$** | | | |
| $E_a, P_a$ res. | 0.22 (-1.14,1.49) | 10109.01 | 1.00 |
| $E_a, P_a$ immi. | 1 (-0.11,2.17) | 14580.06 | 1.00 |
| $E_a, P_s$ res. | 0.14 (-1.05,1.2) | 9189.89 | 1.00 |
| $E_a, P_s$ immi. | -0.04 (-1.46,1.38) | 25623.98 | 1.00 |
| $E_s, P_a$ res. | 0.24 (-1.31,1.71) | 19833.64 | 1.00 |
| $E_s, P_a$ immi. | -0.49 (-1.9,0.92) | 20427.56 | 1.00 |
| $E_s, P_s$ res. | -0.25 (-1.54,1) | 16405.30 | 1.00 |
| $E_s, P_s$ immi. | -0.36 (-1.67,0.94) | 18126.53 | 1.00 |
| **New associate bias $\beta_n$** | | | |
| $E_a, P_a$ res. | 0.3 (-1.19,1.8) | 23100.79 | 1.00 |
| $E_a, P_a$ immi. | 0.43 (-1.04,1.91) | 24623.23 | 1.00 |
| $E_a, P_s$ res. | 0.65 (-0.69,1.81) | 5928.67 | 1.00 |
| $E_a, P_s$ immi. | -0.71 (-2.23,0.85) | 15223.34 | 1.00 |
| $E_s, P_a$ res. | -0.09 (-1.64,1.53) | 27264.13 | 1.00 |
| $E_s, P_a$ immi. | -0.13 (-1.69,1.45) | 26157.39 | 1.00 |
| $E_s, P_s$ res. | 0.41 (-1.36,2.21) | 2827.70 | 1.00 |
| $E_s, P_s$ immi. | 0.24 (-1.19,1.64) | 9807.23 | 1.00 |
| **Behavioral conservatism $\alpha$** | | | |
| | 1.46 (1.08,2) | 4551.06 | 1.00 |
| **SD individals** | | | |
| rho (logit scale) | 2.43 (1.59,3.47) | 4635.11 | 1.00 |
| sigma (logit scale) | 7.01 (4.97,9.66) | 4996.06 | 1.00 |
| $\beta_p$ | 1.04 (0.74,1.43) | 3172.60 | 1.00 |
| $\beta_n$ | 1.08 (0.1,2.39) | 4230.11 | 1.00 |
| alpha (log scale) | 1.92 (0.17,4.14) | 1450.48 | 1.00 |

Table S7: **Summary of model estimates, SL3.** Table contains mean values (89% HDPI), number of effective samples, and Gelman's Rhat. Parameter estimates and SD of varying effects across individuals are reported. The data underlying this table can be found in our data and code repositoryhttps://doi.org/10.17617/3.FXC12W.

| condition | value | n. eff. | Rhat |
|---|---|---|---|
| **Recent information bias $\rho$** | | | |
| $E_a, P_a$ res. | 0.3 (0.13,0.57) | 5949.42 | 1.00 |
| $E_a, P_a$ immi. | 0.2 (0.09,0.42) | 3406.64 | 1.00 |
| $E_a, P_s$ res. | 0.32 (0.16,0.56) | 4761.78 | 1.00 |
| $E_a, P_s$ immi. | 0.23 (0.1,0.46) | 5039.94 | 1.00 |
| $E_s, P_a$ res. | 0.42 (0.17,0.72) | 10619.61 | 1.00 |
| $E_s, P_a$ immi. | 0.23 (0.08,0.52) | 4443.83 | 1.00 |
| $E_s, P_s$ res. | 0.25 (0.11,0.49) | 6560.86 | 1.00 |
| $E_s, P_s$ immi. | 0.12 (0.04,0.31) | 5259.67 | 1.00 |
| **Social information bias $\sigma$** | | | |
| $E_a, P_a$ res. | 0.26 (0.07,0.62) | 18099.07 | 1.00 |
| $E_a, P_a$ immi. | 0.49 (0.18,0.81) | 12818.45 | 1.00 |
| $E_a, P_s$ res. | 0.18 (0.05,0.46) | 14287.71 | 1.00 |
| $E_a, P_s$ immi. | 0.22 (0.06,0.54) | 16235.82 | 1.00 |
| $E_s, P_a$ res. | 0.24 (0.07,0.6) | 21053.66 | 1.00 |
| $E_s, P_a$ immi. | 0.32 (0.09,0.7) | 15930.05 | 1.00 |
| $E_s, P_s$ res. | 0.16 (0.04,0.46) | 17341.29 | 1.00 |
| $E_s, P_s$ immi. | 0.29 (0.09,0.62) | 14814.41 | 1.00 |
| **$\sigma$ slope** | | | |
| $E_a, P_a$ res. | -0.81 (-2.14,0.43) | 2905.82 | 1.00 |
| $E_a, P_a$ immi. | -1.43 (-2.55,-0.37) | 5449.14 | 1.00 |
| $E_a, P_s$ res. | -1.13 (-2.26,-0.28) | 5043.10 | 1.00 |
| $E_a, P_s$ immi. | -1.56 (-2.62,-0.6) | 10798.47 | 1.00 |
| $E_s, P_a$ res. | -0.4 (-1.5,0.64) | 7611.25 | 1.00 |
| $E_s, P_a$ immi. | -0.55 (-1.71,0.62) | 2768.85 | 1.00 |
| $E_s, P_s$ res. | -1.55 (-2.8,-0.39) | 9572.58 | 1.00 |
| $E_s, P_s$ immi. | -0.83 (-1.76,0.03) | 5005.61 | 1.00 |
| **Payoff bias $\beta_p$** | | | |
| $E_a, P_a$ res. | 0 (-1.35,1.26) | 5562.69 | 1.00 |
| $E_a, P_a$ immi. | 0.75 (-0.25,1.87) | 12397.06 | 1.00 |
| $E_a, P_s$ res. | 0.17 (-0.94,1.13) | 9134.40 | 1.00 |
| $E_a, P_s$ immi. | -0.17 (-1.53,1.24) | 22828.06 | 1.00 |
| $E_s, P_a$ res. | 0.08 (-1.42,1.53) | 13039.27 | 1.00 |
| $E_s, P_a$ immi. | -0.69 (-2.03,0.73) | 9406.36 | 1.00 |
| $E_s, P_s$ res. | -0.34 (-1.73,1.02) | 19267.03 | 1.00 |
| $E_s, P_s$ immi. | -0.11 (-1.32,1.09) | 15553.20 | 1.00 |
| **New associate bias $\beta_n$** | | | |
| $E_a, P_a$ res. | 0.34 (-1.17,1.87) | 17681.93 | 1.00 |
| $E_a, P_a$ immi. | 0.07 (-1.4,1.59) | 20084.64 | 1.00 |
| $E_a, P_s$ res. | 1.02 (-0.03,1.96) | 10361.35 | 1.00 |
| $E_a, P_s$ immi. | -0.23 (-1.49,0.99) | 15779.76 | 1.00 |
| $E_s, P_a$ res. | -0.19 (-1.84,1.42) | 22523.01 | 1.00 |
| $E_s, P_a$ immi. | -0.2 (-1.78,1.37) | 19424.41 | 1.00 |
| $E_s, P_s$ res. | 0.94 (-0.61,2.44) | 9078.02 | 1.00 |
| $E_s, P_s$ immi. | 0.85 (-0.32,2.04) | 11727.84 | 1.00 |
| **Behavioral conservatism $\alpha$** | | | |
| | 1.76 (1.34,2.32) | 2286.51 | 1.00 |
| **SD individals** | | | |
| rho (logit scale) | 1.67 (0.95,2.54) | 2025.25 | 1.00 |
| sigma (logit scale) | 4.71 (2.92,6.86) | 4158.91 | 1.00 |
| sigma slope (logit scale) | 0.89 (0.66,1.18) | 4501.67 | 1.00 |
| $\beta_p$ | 0.92 (0.08,2.14) | 3328.69 | 1.00 |
| $\beta_n$ | 0.72 (0.05,1.96) | 3377.31 | 1.00 |
| alpha (log scale) | 1 (0.09,2.52) | 1087.27 | 1.00 |
